# Immune profiling in M. tuberculosis infection enables stratification of patients with active disease

**DOI:** 10.1101/581298

**Authors:** Darragh Duffy, Elisa Nemes, Alba Llibre, Vincent Rouilly, Elizabeth Filander, Hadn Africa, Simbarashe Mabwe, Lungisa Jaxa, Bruno Charbit, Munyaradzi Musvosvi, Humphrey Mulenga, the *Milieu Intérieur* Consortium, Stephanie Thomas, Mark Hatherill, Nicole Bilek, Thomas J Scriba, Matthew L Albert

## Abstract

Tuberculosis (TB) is caused by *Mycobacterium tuberculosis* (*Mtb*) infection and is a major public health problem with an estimated 1.7 billion persons infected worldwide. Clinical challenges in TB include the lack of a blood-based test for active disease, and the absence of prognostic biomarkers for early treatment response. Current blood based tests, such as QuantiFERON-TB Gold (QFT), are based on an IFNγ readout following *Mtb* antigen stimulation. However, they do not distinguish active TB disease from asymptomatic *Mtb* infection. We hypothesized that the use of TruCulture, an improved immunomonitoring method for whole blood collection and immune stimulation, could improve the discrimination of active disease from latent *Mtb* infection. To test our hypothesis, we stimulated whole blood from active TB patients (before and after successful treatment), comparing them to asymptomatic latently infected individuals. *Mtb*-specific antigens (ESAT-6, CFP-10, TB7.7) and live bacillus Calmette-Guerin (BCG) were used for TruCulture stimulation conditions, with direct comparison to QFT. Protein analyses were performed on the culture supernatants using ELISA and Luminex multi-analyte profiling. TruCulture showed an ability to discriminate active TB cases from latent controls (p < 0.0001, AUC = 0.81, 95% CI: 0.69-0.93) as compared to QFT (p = 0.47 AUC = 0.56, 95% CI: 0.40-0.72), based on an IFNγ readout after *Mtb* antigen stimulation. The stratification of the two groups could be further improved by using the *Mtb* Ag/BCG IFNγ ratio response (p < 0.0001, AUC = 0.918, 95% CI: 0.84-0.98). We also identified additional cytokines that distinguished latent infection from TB disease; and show that the primary differences between the TruCulture and QFT systems were a result of higher levels of non-specific innate immune activation in QFT tubes, due to the lack of a buffering solution in the latter. We conclude that TruCulture offers a next-generation solution for whole blood stimulation and immunomonitoring with the possibility to discriminate active and latently infected persons.

## Introduction

Tuberculosis (TB) is a global public health problem, with an estimated 1.7 billion persons latently infected by *Mycobacterium tuberculosis* (*Mtb*)^1,2^. Most exposed individuals mount an effective immune response that controls infection, however the host response generally fails to clear the bacteria resulting in a clinically asymptomatic state^3^. An estimated 10% of subjects with chronic infection progress to active disease at some point in their life, translating into approximately 10 million progressing to TB disease annually^3^.

Treatment regimens can achieve clearance of *Mtb* and infected persons in non-endemic regions are typically recommended for therapy based on the diagnosis of infection. By contrast, due to the burden of infected persons in endemic regions, strategies typically prioritize patients with active disease for treatment with the goal to limit transmission. As a result, there is a critical need to diagnose active disease and to distinguish it from latent infection. Diagnosis of active TB disease can be achieved by microscopy, PCR or culture-based detection of *Mtb* presence in sputum. However, some TB patients cannot produce sputum and in general, it would be preferable to utilize blood based clinical assays, but available methods do not support stratification of patients with active disease from those with latent infection. Instead, available whole blood assays are used to distinguish infected from uninfected persons. Specifically, detection of infection is based on stimulation with *Mtb* antigens, followed by an IFNγ release assay (IGRA), such as the QuantiFERON–TB Gold assay (QFT) or T-SPOT.TB assay. QFT utilizes *Mtb-specific* antigens ESAT-6, CFP-10, and TB7.7 to stimulate immune cells in a blood collection tube with IFNγ secretion readout by ELISA, and T-SPOT.TB uses similar antigens with IFNγ production captured by an ELISPOT assay. Both tests offer clear advantages as compared to tuberculin skin tests (TST), which requires intradermal injection and produces false positive results in BCG-vaccinated individuals^4^.

We hypothesized that an improved method for whole blood collection and immune stimulation could achieve discrimination of active disease from latent *Mtb* infection. Sources of technical variability with QFT include the broad range of blood volumes collected (0.8-1.2mL) and the flexible incubation times indicated in the manufacturer’s protocol (16-24 hours) and recent studies have addressed these sources of variance^4,5^. Despite these improvements QFT has limitations for use as a diagnostic or screening tool in TB endemic countries, but is used to monitor immune responses in vaccine clinical studies and treatment trials^6^.

We have previously described the use of TruCulture^®^ (TruC) devices, a syringe based whole blood collection and incubation system that allows immunomonitoring in response to diverse immune agonists, quantified using proteomic^7^ or transcriptional^8,9^ assays. Specifically we have demonstrated greater reproducibility in multi-centre studies, as compared to conventional PBMC stimulation^10^. Notably, TruC showed significantly lower non-specific cytokine levels in the unstimulated control (Null) tube as compared to PBMC stimulation^10^. This resulted in improved reproducibility across different centres, and greater signal-to-noise ratios for induced immune responses. Given these findings in healthy donors, we aimed to evaluate if the TruC sampling and stimulation method is applicable for the immunomonitoring of TB patients. As shown herein, we demonstrated the ability to more accurately classify patients with active disease and latently infected persons using TruC stimulation, as compared to QFT. The absence of perturbed immune responses in successfully treated patients highlighted the potential of this strategy for use in clinical evaluation of therapeutics and vaccine candidates. We suggest that TruC stimulation systems may address a key unmet diagnostic need, with the possibility to support screening programmes and vaccine programs where QFT conversion is being used as a clinical endpoint.

## Methods

### Participant groups

25 healthy adults with asymptomatic, latent *Mtb* infection (LTBI), defined by a positive QuantiFERON-TB Gold In-Tube (QFT+) assay (Qiagen, Germany), and 25 HIV-negative adults with TB disease (TB), defined by a positive sputum XpertMTB/RIF test (Cepheid, United States) were identified and recruited at the South African Tuberculosis Vaccine Initiative (SATVI), Worcester, South Africa^11^. The LTBI control group was age, sex and ethnicity matched to the active TB disease patient group. Blood was collected prior to treatment initiation in active TB cases and again 12-18 months later, after successful completion of treatment and having been declared clinically cured (V2, n=18). For the LTBI controls, blood was also collected at a second time-point, 12-18 months after the initial visit (V2, n=19). TB patient and LTBI control characteristics are described in Table S1. The TB clinical study, protocols and informed consent forms were approved by the Human Research Ethics Committee of the University of Cape Town (ref: 234/2015). Healthy donor blood from a French population in a non-endemic TB setting was obtained from the CoSImmGEn cohort of the Investigation Clinique et Accès aux Ressources Biologiques (ICAReB) platform, Centre de Recherche Translationnelle, Institut Pasteur, Paris, France. Written informed consent was obtained from all study participants.

### Whole blood stimulations

TruC tubes (Myriad RBM) were batch prepared and maintained at −20°C until time of use. To prepare TruC TB antigen tubes, 3 QFT TB antigen tubes (the QFT Gold system was used, as the study was performed prior to introduction of the QFT Gold Plus) were rinsed with 2mL of TruC media and the media transferred into empty TruC tubes to maintain the same concentration of *Mtb* antigens as found in QFT. Live bacillus Calmette-Guerin (BCG; Connaught strain, Sanofi Pasteur) tubes were prepared to have a final concentration of 10^5^ bacteria/mL. 1mL of whole blood was collected directly in QFT Gold tubes (Nil, TB Antigens, Mitogen) (Qiagen) according to manufacturers’ instructions. Whole blood was also collected in Sodium Heparin tubes and 1mL transferred into TruC tubes (total volume 3mL). Unstimulated control tubes from both stimulation systems are referred to as Null throughout to avoid confusion. QFT and TruC tubes were processed within 30minutes of blood draw, inserted into a dry block incubator, and maintained at 37°C (±1°C) room air for 22 hrs (+/− 15 minutes) as previously described^7^. At the end of the incubation period, QFT tubes were centrifuged and TruC tubes were opened and a valve was inserted to separate the sedimented cells from the supernatant, stopping the stimulation reaction. Separate supernatant aliquots were prepared for ELISA and Luminex testing and frozen at −80°C until analysis.

### Multi-analyte protein profiling

Supernatants from QFT and TruC tubes were analyzed for IFNγ by standard ELISA (Qiagen) and values were expressed in IU/mL, calculated by subtraction of values from the relevant nonstimulated controls. Luminex xMAP technology was used to measure 32 proteins in the same samples (Myriad RBM). Samples were measured according to CLIA guidelines (set forth by the USA Clinical and Laboratory Standards Institute). The 32 measured analytes were organized on 3 multiplex arrays, and a single batch of reagents was used for testing all samples per timepoint. The least detectable dose (LDD) for each assay was derived by averaging the values obtained from 200 runs with the matrix diluent and adding 3 standard deviations to the mean. The lower limit of quantification (LLOQ) was determined based on the standard curve for each assay and is the lowest concentration of an analyte in a sample that can be reliably detected and at which the total error meets CLIA requirements for laboratory accuracy. The lower assay limit (LAL) is the lowest value read out after application of the standard curve and use of curve-fitting algorithms. In most instances, the LAL is less than the LDD and the LLOQ. To enable a direct comparison between both stimulation systems protein concentrations were calculated to pg/ml of whole blood, which integrated the original dilution factors.

### Statistical analysis

Normality of induced cytokine responses were evaluated (Shapiro Wilk test), and accordingly, T tests were performed for two group comparisons with Qlucore Omics Explorer, v.3.4(11) (Qlucore) or GraphPAD Prism (version 6). For comparisons between pre and post treatment paired *t* tests were performed with GraphPAD Prism. Multiple testing correction was performed and false discovery rate (FDR)-adjusted p values (referred to herein as q values) are reported. Sample size calculation was performed and 25 persons per group was chosen to have sufficient power to detect differences due to TB disease. Calculations were based on previously published induced immune responses in *Mtb* infected vs. healthy controls^5^. For the analysis of donors in a non-endemic setting, non-parametric ANOVA tests were performed due to the smaller sample size (n= 10). Dot plot graphs were compiled with GraphPad Prism and heat maps were generated using Qlucore v.3.4(11). Receiver operating characteristic (ROC) curves were calculated and compared with R Studio v.3.3.1 pROC package and results drawn with graphical package ggplot2 v.2.1.0.

## Results

### Improved discrimination of patient groups using TruCulture TB Ag stimulation

To enable direct comparison between TruC and QFT, we transferred *Mtb* antigens from QFT into TruC tubes as described (see Methods). All other tubes were prepared as previously described^7^. We sampled blood from active TB patients and LTBI persons (see Methods), and measured induced IFNγ production utilizing ELISA. Confirming previous reports^12^, QFT assays did not stratify TB and LTBI groups (Fig 1a, p=0.47). By contrast, the TruC method, using the same *Mtb* antigens and IFNγ readout, showed a significantly higher stimulation index in TB patients as compared to LTBI controls (p = 0.001, Fig 1b). Inclusion criteria for defining LTBI cases was based on historical QFT positivity (IFNγ concentrations above 0.35 IU/mL), confirmed upon re-testing (Fig 1a). Indicating distinct parameterization between the two assays, when this pre-defined cut off was applied to the TruC results, only 9 LTBI cases and 17 TB patients scored positive (Fig 1b).

**Figure 1.**
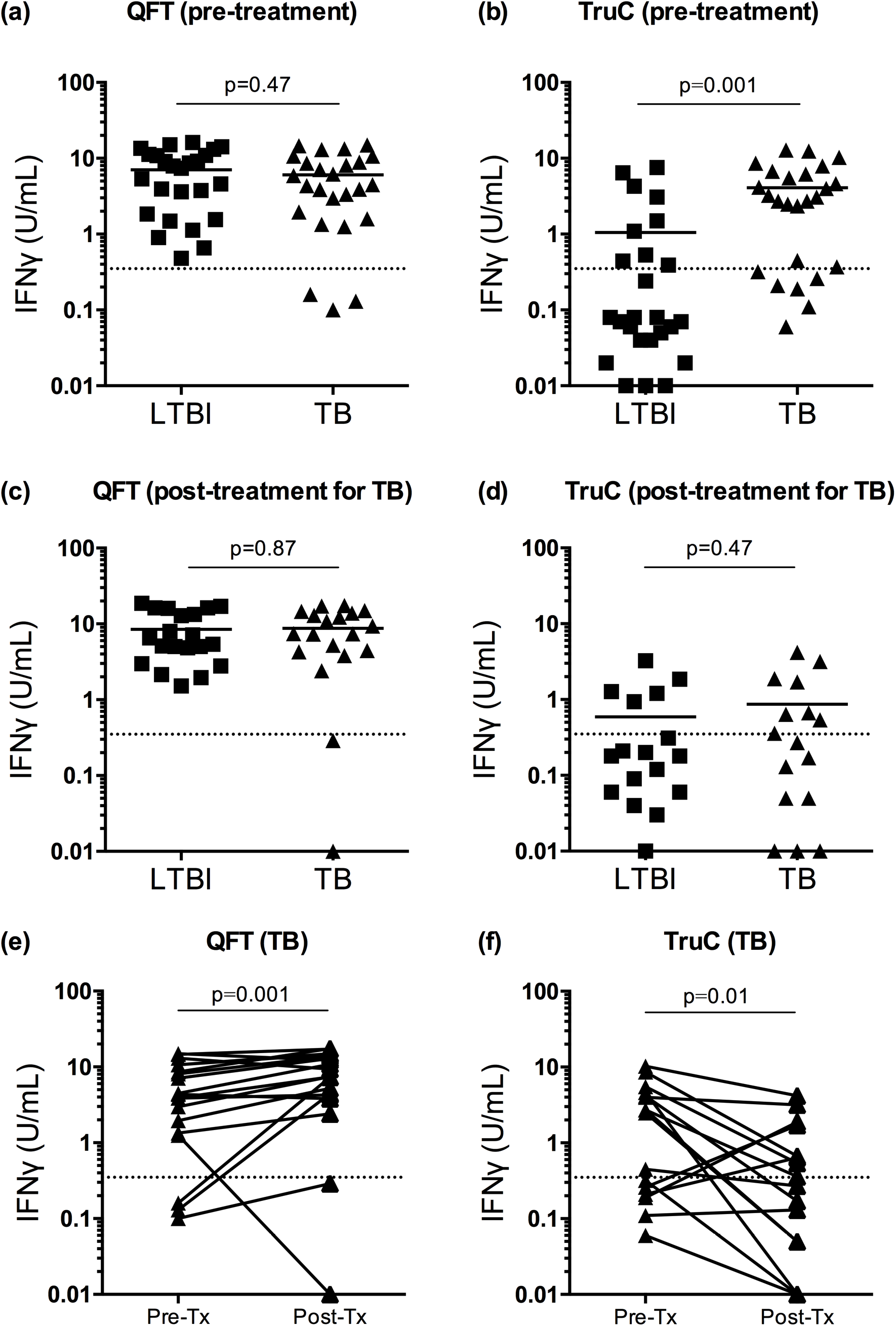
IFNγ *Mtb* Ag response. IFNγ response following *Mtb* Ag stimulation and subtraction of the Null control in LTBI and TB patients in (a) QFT tubes pre-treatment, (b) TruC tubes pre-treatment, (c) QFT tubes after successful antibiotic treatment in TB patients, (d) TruC tubes after successful antibiotic treatment in TB patients. (a, b n=25/25; c,d n=19/18 latent/active, Student T test), bars represent the mean values, dotted line is QFT positive cut off 0.35 IU/ml). Paired IFNγ responses following *Mtb* Ag stimulation and subtraction of the Null control in TB patients pre and post treatment in (e) QFT tubes and (f) TruC tubes (Paired T test).

All patients with active disease were treated, and 18 agreed to retesting after 12-18 months and following successful treatment. We also retested 19 LTBI controls after a similar 12-18 month time interval. At this time point no differences were observed between the LTBI and treated TB patients with either QFT or TruC systems (Fig 1c & d). When the effect of treatment on TB patients was directly examined, both QFT and TruC assays showed significant differences (pre-vs. post-treatment, paired T test, Fig 1e & f). Paradoxically, patients showed an increased IFNγ response in QFT (p=0.001) when comparing post-vs. pre-treatment cytokine levels; whereas the majority of patients showed the expected decrease in IFNγ responses as measured by TruC assays (p =0.01).

### Multiple cytokine responses stratify active and latent TB after TruC Mtb Ag stimulation

To assess the value of measuring additional inflammatory cytokines, we performed CLIA certified Luminex multi-analyte profiling on all supernatants, quantifying a total of 32 cytokines, chemokines and growth factors. Statistical analysis identified 7 proteins that were differentially expressed between TB and LTBI groups (q < 0.01, Fig 2a, c, Table S2), in the *Mtb* antigen TruC supernatants, whereas no differences were observed in the respective QFT assays (Fig 2b, d, Table S2). A heat map representation of the TruC results illustrates 5 proteins that showed higher responses (IFNγ, IL-18, IL-1RA, IL-8, CCL4), and 2 with lower responses (CCL11, Factor VII) in active TB as compared to LTBI patients (Fig 2a), with no discernible pattern observed in the QFT stimulations (Fig 2b). Individual plots of protein concentration are also depicted (Fig 2c & d). Following successful treatment of the TB group there were no significant differences between the treated TB patients and LTBI controls (Fig 2e). This analysis indicated that TruC stimulation could reveal multiple immune perturbations in patients with active TB disease.

**Figure 2.**
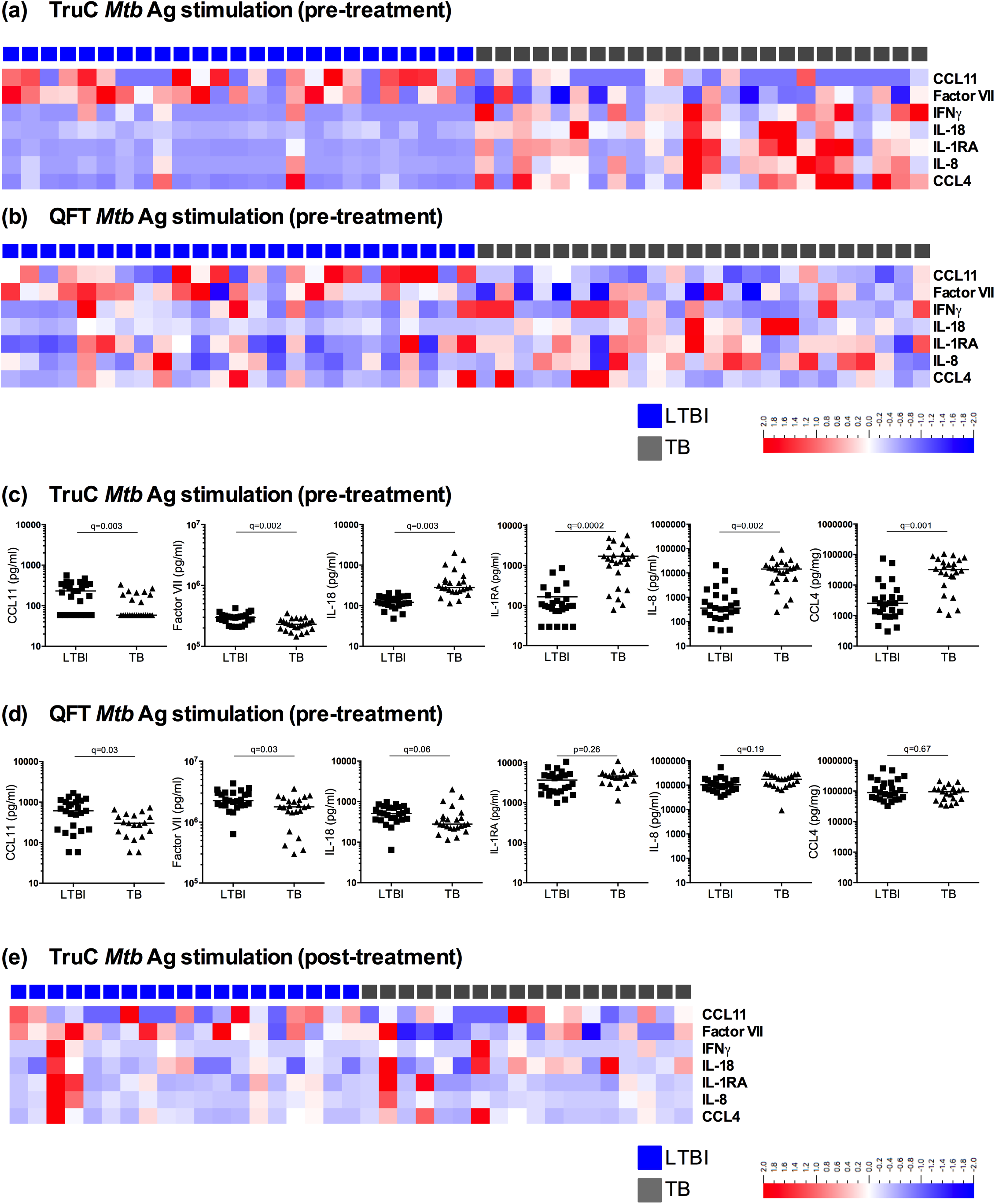
Differential cytokine responses in *Mtb* infection versus TB disease. Heat maps of relative cytokine expression levels segregated by patient group (LTBI: blue, TB: grey) after *Mtb* Ag stimulation in TruC (a) or QFT (b) tubes prior to treatment, and (e) TruC *Mtb* Ag stimulation after successful antibiotic treatment of the TB patient group. (c) Dot plot representations of the cytokine concentrations in TruC (c) or QFT (d) tubes prior to treatment. Cytokines included showed a significant difference (q<0.01) following FDR adjusted T tests between LTBI and TB groups. (a, b, c, d n=25/25; e n=19/18 latent/active, bars represent the mean values, q value: FDR corrected Student T test).

### TruC BCG stimulation revealed additional immune response differences and improved patient classification

Given its use as a TB vaccine and its ability to trigger an innate response in whole blood^13^, we included BCG as an additional TruC stimulation condition. Of the 15 proteins that were induced by BCG, 11 were differentially expressed (q < 0.01) between the two patient groups (Fig 3a, Table S3). Interestingly, the BCG induced proteins were higher in the LTBI group, with 4 of the most differentially expressed proteins being IL-1 family members (IL-1α, IL-1β, IL-1RA, and IL-18) (Fig 3b). Interestingly, IFNγ was nominally significant, higher in the LTBI group (p = 0.04) (Fig 3b). Again, the immune responses in TB patients post-treatment, normalized to those seen in LTBI controls (Fig. S1).

**Figure 3.**
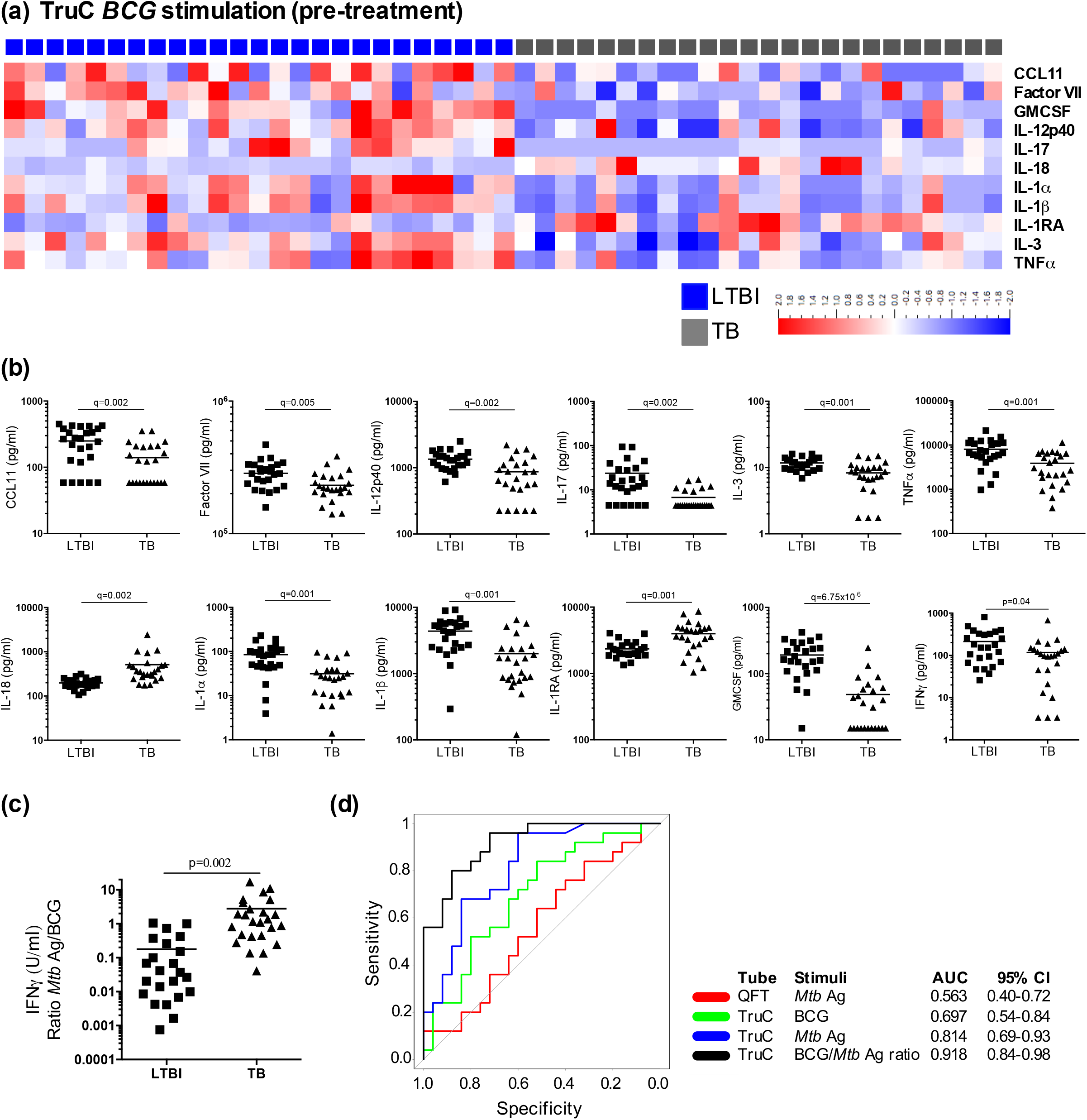
BCG induced immune responses in *Mtb* infection versus TB disease. (a) Heat map of relative cytokine expression levels segregated by patient group (LTBI: blue, TB: grey) after BCG TruC stimulation, and identification of differential proteins after FDR adjusted T tests between LTBI and TB groups. (b) Dot plot representations of the cytokine concentrations of differential proteins between LTBI and TB groups: CCL11, Factor VII, IL-12p40, IL-17, IL-3, TNFα, IL-18, IL-1α, IL-1β, IL-1RA, GMCSF. (c) IFNγ ratio (U/ml) of *Mtb* Ag/BCG stimulation for LTBI and TB patients. (d) ROC curve analysis of IFNγ ratio to *Mtb* Ag/BCG stimulation (black), IFNγ concentrations of TruC *Mtb* Ag (blue), TruC BCG (green), and QFT *Mtb* Ag (red) stimulations. (n=25/25, bars represent the mean values, q value: FDR corrected Student T test).

Given that the pattern of stimulation was inverse to that observed using *Mtb* antigen (ie higher in LTBI compared to TB), we predicted that BCG induced cytokines could be leveraged for improving the stratification of patient groups. We therefore calculated a composite index, the ratio of *Mtb* Ag and BCG induced IFNγ response, which showed a >10-fold difference between the two patient groups (p=0.002, Fig 3c), and a ROC area under the curve (AUC) of 0.918 (95% CI: 0.84-0.98) (Fig 3d). Notably, this AUC was superior than those achieved for the individual tests: TruC *Mtb* Ag (AUC 0.814, 95% CI: 0.69-0.93), or BCG (AUC 0.697, 95% CI: 0.54-0.84); and the QFT *Mtb* Ag (AUC 0.563, 95% CI: 0.40-0.72), and a bootstrap test for correlated ROC curves revealed statistically significant improvements; TruC *Mtb* Ag, p = 0.02; QFT *Mtb* Ag p < 0.0001. While TruC *Mtb* Ag alone performed better than QFT *Mtb* Ag (p = 0.04), it was not superior to BCG TruC (p = 0.11). These findings demonstrate the potential advantage of combining peptide antigen and complex stimuli for improved patient classification (Fig 3d).

### QFT negative control tubes have high non-specific cytokine activation

To examine the underlying differences between TruC and QFT, we considered the non-specific activation using the respective Null control conditions. To avoid potential artefacts caused by outlier measurements, we performed pre-filtering based on variance (σ/σ_max_ = 3.25×10^−5^), which led to removal of 9 proteins that showed low variance across all conditions. Statistical testing on the remaining 23 proteins measured revealed highly significant differences (null conditions QFT vs. TruC q < 0.01) with all proteins showing higher concentrations in the QFT tubes (Fig 4a, b, Table S4). IL-6, IL-1β and CCL2 were the 3 most differentially expressed proteins (Fig 4c). Notably these differences were independent of disease status, as all proteins remained significantly different after regressing for patient status (TB or LTBI).

**Figure 4.**
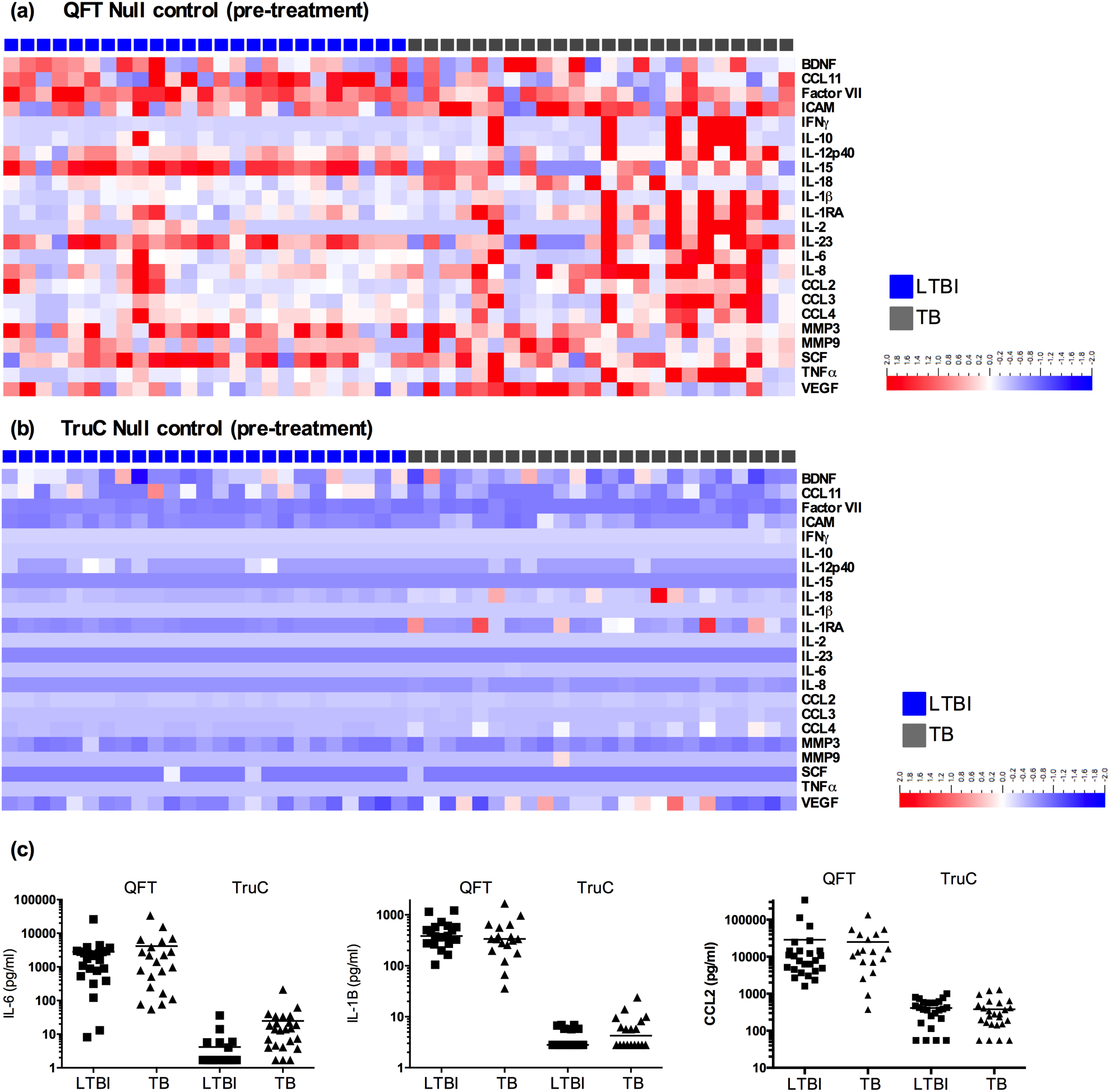
Differential cytokines in QFT and TruC Null tubes. Heat maps of relative cytokine expression levels segregated by patient group (LTBI: blue, TB: grey) in (a) QFT and (b) TruC Null tubes for 22 out of 32 cytokines measured, selected based on variance (σ/σ_max_ = 0.138). (c) Concentrations of IL-6, IL-1β, CCL2 in QFT and TruC Null tubes in LTBI and TB patients. (n=25/25, bars represent the mean values).

To further validate this analysis, we performed additional experiments in healthy uninfected European donors. The same QFT and TruC stimulations were performed as described above. Additionally, we investigated the hypothesis that the QFT tube or the TruC media might account for the observed variability between the Null conditions. We tested conditions in which blood collection was performed in the TruC tubes followed by transfer of the blood/media mixture into a QFT Null tube; as well as the converse, initial blood collection and mixing in QFT tubes followed by transfer into a TruC Null tube containing media in the absence of additional stimuli. A comparison between QFT and TruC, negative control tubes and *Mtb* Ag tubes, showed similar results to the TB patients and LTBI controls, with significantly higher levels of innate cytokines in QFT (examples of IL-6, IL-1β and CCL2 shown for comparison with prior results illustrated in Fig 5a-c). Strikingly, in both of the tube transfer conditions, the cytokine levels reflected the TruC condition and indicated that the presence of the TruC media minimized the non-specific innate cell activation observed when using the QFT tubes. Unexpectedly, 1 donor showed elevated IFNγ responses in both stimulation systems (Fig 5d, highlighted in blue), however the fold change of the *Mtb* Ag over the Null response was 4-fold in QFT, and 16-fold in TruC, illustrating the improved signal-to-noise achievable for induced antigen specific immune responses using TruC (Fig 5d). This particular donor was also an outlier for other cytokine responses (e.g., IL-1β, IL-6) in *Mtb* Ag and BCG TruC stimulations (Fig 5a-c, highlighted in blue). These combined results removed possible confounding factors due to TB infection and demonstrated that TruC media and the conditions reported facilitate an improved method for immune stimulation and immune monitoring. We conclude that the use of TruC may provide considerable advantages if further developed as a method for immunmonitoring in TB clinical studies and patient management strategies.

**Figure 5.**
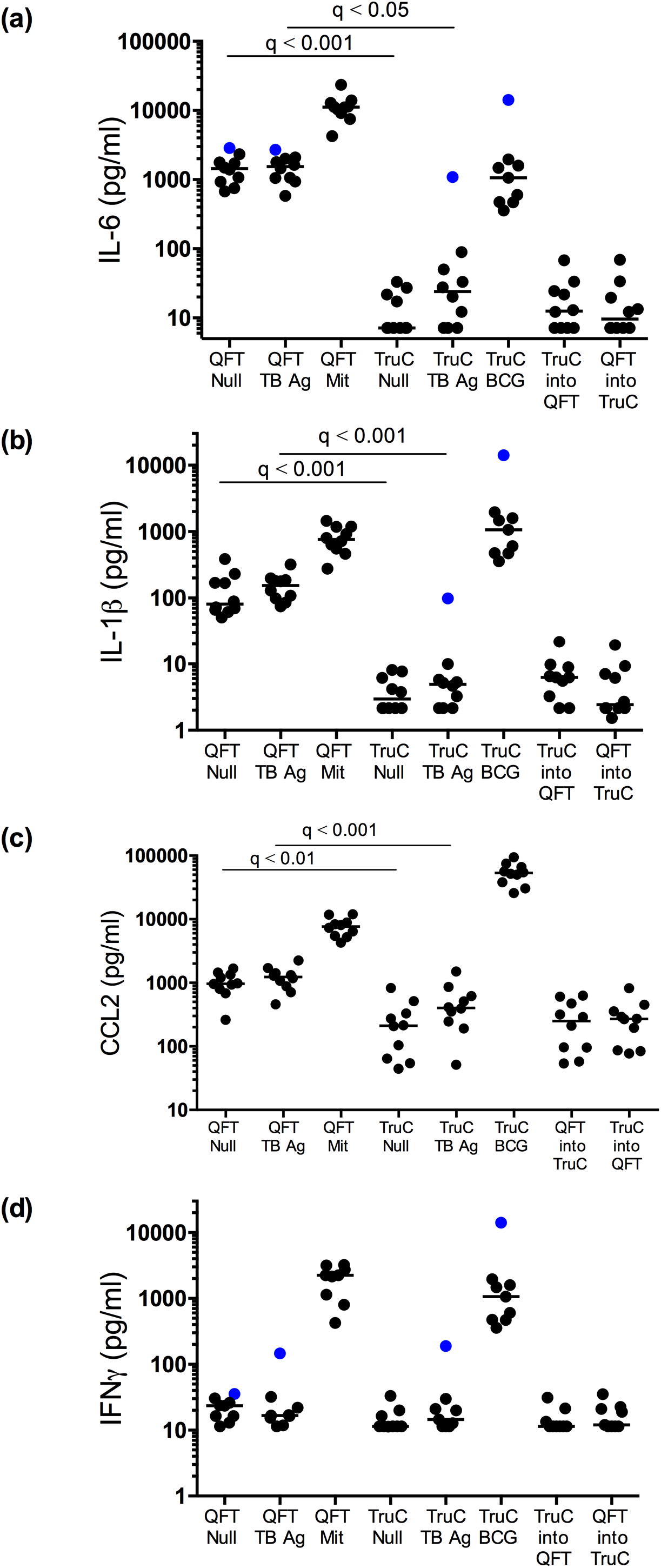
Cytokine responses in donors from a non-endemic TB region. Concentrations of (a) IL-6, (b) IL-1β, (c) CCL2, and (d) IFNγ in QFT Null, QFT TB Ag, QFT Mitogen, TruC Null, TruC TB Ag, TruC BCG, and mixed cultures of TruC-QFT and QFT-TruC Null conditions in healthy donors from a non-endemic region (n=10, bar represents the median values, Friedman test with Dunn’s multiple comparisons test, outlier donor in blue).

## Discussion

Blood-based immunomonitoring is increasingly used in clinical studies due to the ease of sampling and the possibility of longitudinal measurements during medical interventions. TB is a highly relevant example of how such an approach can be applied to monitor functional immune responses in routine clinical applications, and this approach has been extended to cytomegalovirus infection and transplantation settings^14^. However, the use of QFT blood-based tests in TB endemic countries has been limited by their poor ability to discriminate active TB disease from asymptomatic infection. Such stratification is key for TB control strategies that are focused on treating infection to reduce risk for re-activation, thus diminishing the chance for new transmissions.

We demonstrated here a clear advantage of utilizing an alternative immunomonitoring tool, TruCulture, for the analysis of induced immune responses in TB disease. Utilizing the same *Mtb* antigens and IFNγ readout as the QFT assay, TruC showed significant differential induced IFNγ expression in patients with active disease as compared to LTBI, differences that have not been achieved using the QFT test^14^. Furthermore, stimulation with live BCG yielded a unique signature, with higher cytokine expression found in LTBI as compared with persons with active disease. We suggest that this pattern of expression is due to the particulate nature of the BCG antigen and the requirement for phagocytosis to achieve antigen processing and presentation^15^, as compared to free peptides. Notably, it has been demonstrated that inflammation limits the ability of antigen presenting cells to capture and present particulate antigen^16^, thus providing a possible explanation for our findings. As a consequence of the distinct pattern of expression, it was possible to combine the *Mtb* Ag and BCG-induced responses in order to improve the classification of active *versus* LTBI individuals. Use of TruC also revealed differential induction of other cytokines, representing both innate and adaptive immune responses. Importantly, we show that such immune response differences may be obscured in the QFT cultures by cytokines that are activated non-specifically, presumably from the activation of myeloid cells in the absence of the TruC liquid media.

The high concentrations of multiple cytokine responses in the non-stimulated control QFT tube was somewhat unexpected. Interestingly, the elevated non-specific immune responses were reminiscent of a previously reported non-specific activation of myeloid cells^10^. To minimize such issues in clinical applications of QFT, as well as in the vast majority of research studies, decision making is restricted to the study of IFNγ responses, with the value of the non-stimulated control being subtracted from the value of the *Mtb* Ag stimulation^5^. While this delivers meaningful information about antigen-specific adaptive responses, our study illustrates the added value of reducing the overall background biological noise due to the method of stimulation^17^, thus supporting improved signal discrimination. In doing so, the signal-to-noise is improved, revealing previously unappreciated differences in antigen-specific signals between active TB and asymptomatic infection, which if further developed as a diagnostic tool could help lessen the so-called IGRA zone of uncertainty^5^. Improving the sensitivity and specificity of tests in these particular individuals could have a considerable impact on clinical decision making^18^, and helping to deliver preventive therapy to those with latent *Mtb* infection or sparing those who do not require prophylaxis from lengthy antibiotic regimens.

Prior studies have reported multi-analyte profiling using Luminex technology. In such studies, QFT stimulations were used and in some instances results could be compared to our findings^19,20^. The ranges reported and those observed in our study were of a similar magnitude (e.g. for IL-1RA, CXCL10^19^, IL-1α, IL1β, IL-2, IL-6, CCL2, CCL3, TNFα^21^). Notable differences were observed in the levels of IL-8 and CCL4, which were an order of magnitude lower than those observed in our study. This may be explained by differences in antibodies used or in the populations studied. Supporting the latter explanation, we observed differences between the South African and French donors, the former group showing higher levels of cytokines in the unstimulated QFT condition. This may reflect: immune cell composition differences and/or immune response variability between populations of distinct genetic ancestry^22^ that may be further amplified by non-specific myleoid cell activation; underlying differences due to latent *Mtb* infection in donors; or other environmental factors. This requires further study and highlights the challenges of non-standardized biomarker studies across different sampling times and sites, and further highlights the need for reproducible sample preparation and analyte testing.

Caveats of our study include the small sample size and the absence of an independent validation cohort. Despite the modest number of patients, highly statistically significant differences were observed with TruC, an indication of the large effect size observed. This also highlights how robust sampling can facilitate the powering of scientific questions with smaller study sizes. Further validation of our findings are planned with the aim to identify clinical questions that would most benefit from the use of TruC devices. Given the recent advances in *ex vivo* whole blood transcriptomic signatures for diagnosing subclinical or active TB disease^23,24^, the requirement for an incubation step may represent a barrier to clinical translation to near-patient testing. Despite the stated limitations, we believe that there is sufficient justification for testing TruC systems as next generation immunomonitoring tools in TB clinical studies.

In summary, given the numerous challenges still present in the TB field and the critical need for better tools, the availability of novel robust and adaptable immunomonitoring tools may support the many ongoing efforts to combat TB worldwide.

## Acknowledgments

We acknowledge the Bill and Melinda Gates Foundation (OPP1114368) for funding the study and additional support from the French Government’s Investissement d’Avenir Program, Laboratoire d’Excellence “ Milieu Intérieur” Grant ANR-10-LABX-69-01. We acknowledge the UTechS CB and Icareb platforms of the Center for Translational Research, Institut Pasteur. EN is an International Society for Advancement of Cytometry (ISAC) Marylou Ingram Scholar.

---

**Figure S1.**
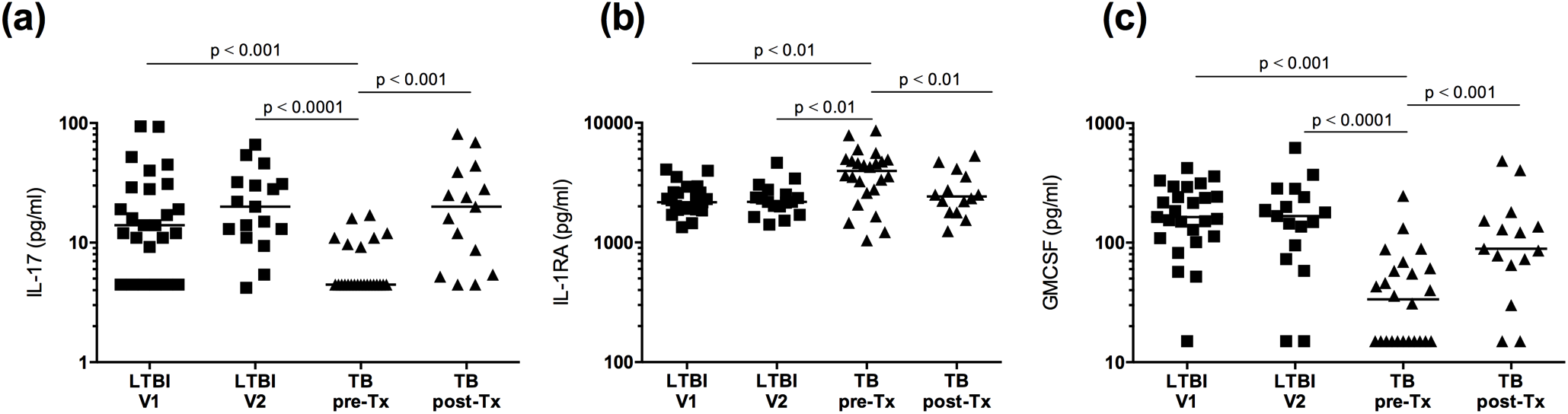
BCG induced immune responses in *Mtb* infection versus TB disease after treatment. Dot plot representations of the cytokine concentrations of differential proteins after BCG stimulation in LTBI (V1 and V2) and TB groups (pre and post-treatment), examples shown for IL-17, IL-1RA, and GMCSF (n=25/25/19/18, Kruskal-Wallis test with Dunn’s multiple comparison test).

**Table S1.**
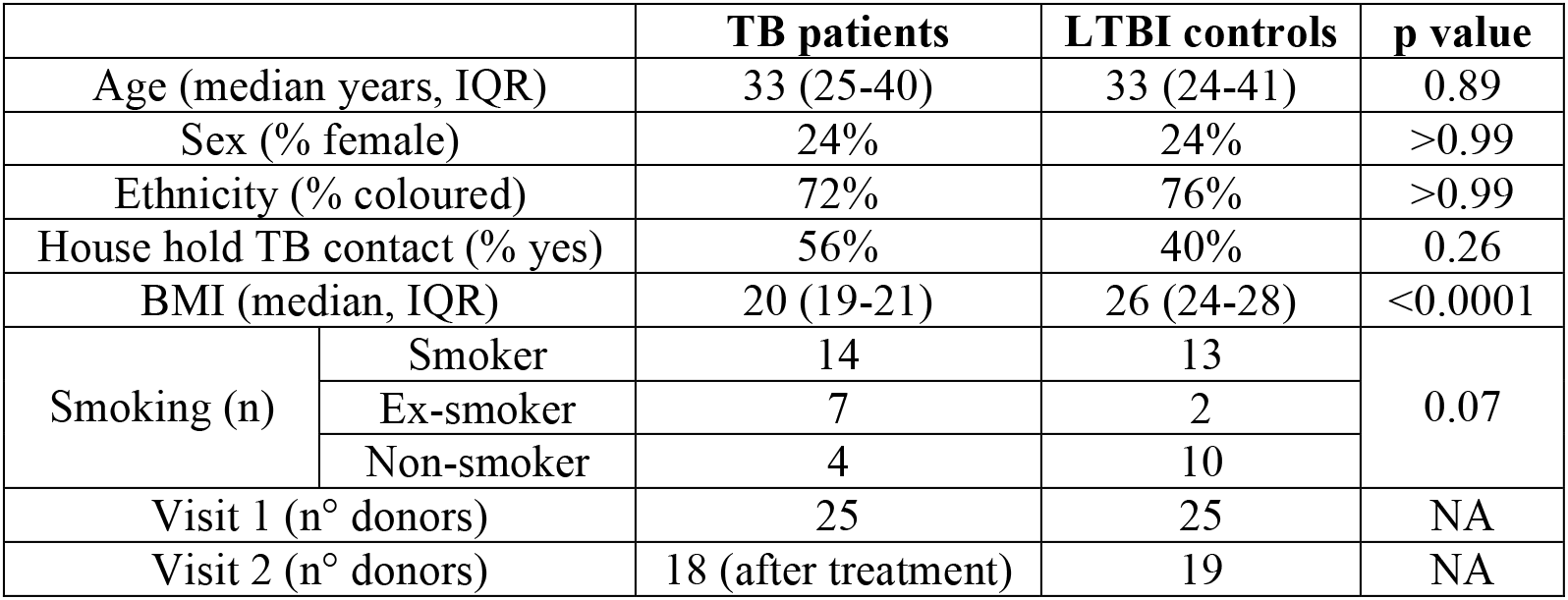
Patient characteristics.

**Table S2.** Q values from ANOVA testing between TB and LTBI groups following *Mtb* Ag stimulation with QFT or TruC tubes, and means with standard deviations (sd) in pg/mL.

| Protein | QFT<br>LTBI v TB<br>(q value) | Mean (sd) pg/mL |  | TruC<br>Latent v<br>Active<br>(q value) | Mean (sd) pg/mL |  |
| --- | --- | --- | --- | --- | --- | --- |
|  |  | LTBI | TB |  | LTBI | TB |
| IL-1RA | 0.26 | 3683<br>(2390) | 4463<br>(2240) | 0.0002 | 165<br>(198) | 1716<br>(1524) |
| CCL4 | 0.67 | 149,384<br>(135,169) | 95709<br>(52797) | 0.001 | 8388<br>(17,310) | 39,059<br>(31,604) |
| IFNγ | 0.7 | 345 (463) | 258 (281) | 0.002 | 21 (41) | 121 (119) |
| IL-8 | 0.19 | 124,060<br>(101,638) | 166,933<br>(85164) | 0.002 | 2047<br>(4593) | 17,412<br>(19,160) |
| Factor<br>VII | 0.03 | 2419040<br>(809,363) | 1,735,739<br>(848,444) | 0.002 | 291,440<br>(58,200) | 232,041<br>(50,946) |
| IL-18 | 0.06 | 532 (226) | 1668 (2146) | 0.003 | 128 (41) | 449 (437) |
| CCL11 | 0.03 | 681 (461) | 327/201 | 0.003 | 240 (148) | 115 (83) |

**Table S3.** Q values from ANOVA testing between TB and LTBI groups following BCG stimulation.

| Protein | LTBI v TB<br>(q value) | TruC mean (sd) pg/ml |  |
| --- | --- | --- | --- |
|  |  | LTBI | TB |
| GMCSF | $6.75 \times 10^{-6}$ | 191 (103) | 48 (52) |
| IL-1α | 0.001 | 84 (56) | 31 (27) |
| IL-1β | 0.001 | 4377 (2201) | 1995 (1713) |
| IL-1RA | 0.001 | 2350 (698) | 3980 (1911) |
| IL-3 | 0.001 | 11 (2) | 8 (3) |
| TNFα | 0.001 | 8047 (4779) | 3887 (2813) |
| IL-12p40 | 0.002 | 1346 (417) | 868 (565) |
| IL-17 | 0.002 | 23 (24) | 6 (3) |
| CCL11 | 0.002 | 249 (136) | 140 (94) |
| IL-18 | 0.002 | 198 (48) | 510 (483) |
| Factor VII | 0.005 | 285,400 (66,662) | 231,625 (57,423) |

**Table S4.** Q values from ANOVA testing between QFT and TruC Null tubes, and means with standard deviations (sd) in pg/mL.

| Protein | QFT v<br>TruC Null<br>(q value) | QFT mean (sd) pg/mL |  | TruC mean (sd) pg/mL |  |
| --- | --- | --- | --- | --- | --- |
|  |  | LTBI | TB | LTBI | TB |
| IL-1β | 5.46 x10<sup>-44</sup> | 481 (320) | 556/535 | 5/2 | 6/4 |
| Factor VII | 9.59 x10<sup>-37</sup> | 2,717,200<br>(884206) | 1,772,739/<br>817,991 | 286,240/<br>63,738 | 231,166/<br>59,543 |
| IL-8 | 3.83x10<sup>-35</sup> | 63,615<br>(39,534) | 116,803<br>(84392) | 121<br>(109) | 2549<br>(3504) |
| IL-10 | 5.54x10<sup>-33</sup> | 49 (135) | 23 (15) | 3 (0) | 3 (1) |
| TNFα | 1.68x10<sup>-31</sup> | 897 (657) | 984 (998) | 10 (4) | 28 (28) |
| IL-6 | 4.90x10<sup>-31</sup> | 2,797 (5,021) | 4,159 (7,673) | 4 (7) | 24 (42) |
| MMP3 | 4.43x10<sup>-30</sup> | 65,624<br>(35,943) | 52,625<br>(28,502) | 9,572 (5,011) | 8,133 (3,677) |
| SCF | 9.66x10<sup>-30</sup> | 570 (232) | 552 (238) | 124 (47) | 116 (24) |
| CCL3 | 8.03x10<sup>-29</sup> | 5,568<br>(3,567) | 10,420<br>(13,736) | 57 (106) | 393 (525) |
| IL1α | 6.58x10<sup>-26</sup> | 15 (9) | 19 (28) | 0 (0) | 0 (0) |
| CCL20 | 6.90x10<sup>-26</sup> | 5,932 (20,978) | 1,344 (1,219) | 79 (216) | 181 (192) |
| IL-12p40 | 1.04x10<sup>-25</sup> | 1,083 (342) | 836 (301) | 339 (198) | 260 (97) |
| IL-23 | 1.69x10<sup>-24</sup> | 4,896 (1,346) | 3,436 (1,685) | 1,600 (0) | 1600 (0) |
| CCL4 | 9.66x10<sup>-21</sup> | 51,716<br>(51,565) | 55,543<br>(53,896) | 904 (585) | 10,807<br>(13,615) |
| IL-1RA | 1.19x10<sup>-16</sup> | 1079 (609) | 1660 (1483) | 88 (42) | 712 (837) |
| CD54 | 2.26x10<sup>-13</sup> | 462,860<br>(275,561) | 777,004<br>(410,222) | 71,600<br>(28,728) | 118,004<br>(74,185) |
| MMP9 | 3.64x10<sup>-11</sup> | 41,060<br>(20,215) | 63,000<br>(84,132) | 16,500 (0) | 17,562<br>(5,205) |
| IL-18 | 3.55x10<sup>-10</sup> | 539 (221) | 1,517 (1,998) | 124 (42) | 433 (492) |
| VEGF | 2.62x10<sup>-8</sup> | 779 (340) | 1,318 (727) | 364 (138) | 530 (309) |
| BDNF | 2.94x10<sup>-7</sup> | 21,040<br>(8,691) | 22,458<br>(13,532) | 13,208<br>(4,624) | 12,183<br>(4,824) |
| CCL11 | 5.66x10<sup>-7</sup> | 794 (543) | 404 (298) | 267 (181) | 143 (95) |
| IFNγ | 1.00x10<sup>-6</sup> | 5 (4) | 11 (11) | 3 (0) | 4 (3) |

